# Spatial transcriptome profiling uncovers metabolic regulation of left-right patterning

**DOI:** 10.1101/2023.04.21.537827

**Authors:** Hisato Yagi, Cheng Cui, Manush Saydmohammed, George Gabriel, Candice Baker, William Devine, Yijen Wu, Jiuann-huey Lin, Marcus Malek, Abha Bais, Stephen Murray, Bruce Aronow, Michael Tsang, Dennis Kostka, Cecilia W. Lo

## Abstract

Left-right patterning disturbance can cause severe birth defects, but it remains least understood of the three body axes. We uncovered an unexpected role for metabolic regulation in left-right patterning. Analysis of the first spatial transcriptome profile of left-right patterning revealed global activation of glycolysis, accompanied by right-sided expression of *Bmp7* and genes regulating insulin growth factor signaling. Cardiomyocyte differentiation was left-biased, which may underlie the specification of heart looping orientation. This is consistent with known *Bmp7* stimulation of glycolysis and glycolysis suppression of cardiomyocyte differentiation. Liver/lung laterality may be specified via similar metabolic regulation of endoderm differentiation. *Myo1d*, found to be left-sided, was shown to regulate gut looping in mice, zebrafish, and human. Together these findings indicate metabolic regulation of left-right patterning. This could underlie high incidence of heterotaxy-related birth defects in maternal diabetes, and the association of PFKP, allosteric enzyme regulating glycolysis, with heterotaxy. This transcriptome dataset will be invaluable for interrogating birth defects involving laterality disturbance.

## Introduction

Left-right (LR) organ asymmetries include evolutionary adaptations enabling animals transitioning from an aquatic environment to survive on land. The most LR asymmetric organ in the body is the cardiovascular system. Its LR asymmetries provide distinct systemic vs. pulmonary circulation required for efficient blood oxygenation with oxygen extraction from air in the lungs. The digestive tract, which initially develops outside the body cavity, undergoes directional gut rotation as it is retracted and compacted into the abdominal cavity, a process also regulated by LR patterning. This allows for lengthening of the intestinal tract required for the increased water absorption essential for survival on land. Hence, disturbance of LR patterning can cause severe birth defects associated with high morbidity and mortality. In patients with disturbance of left-right patterning, such as in heterotaxy with randomization of left-right visceral organ situs, the cardiovascular system is often disrupted, contributing to the high incidence of complex congenital heart defects (CHD) in heterotaxy^1, 2^. Other organ laterality defects also can be observed with heterotaxy, such as gut malrotation that can cause life threatening intestinal volvulus and gut obstruction and ischemia, or biliary atresia with defects in formation of the bile duct that often require liver transplant for long term survival.

The prevention of birth defects involving LR patterning defects will require insights into the mechanism of LR patterning. Unexpected lead were intimated by clinical observations dating back to the 1800’s with Afzelius report of Kartagener’s syndrome with the puzzling triad of defects including complete mirror symmetric positioning of visceral organs (situs inversus totalis), male infertility, and primary ciliary dyskinesia (PCD) in which sinopulmonary disease arises from mucociliary clearance deficiency^3^ due to immotile cilia in the airway. We now understand the association of PCD with both laterality defects and male infertility reflects the common role of motile cilia in LR patterning, airway mucociliary clearance and sperm motility^4^. Also, puzzling is the well described association of maternal diabetes with birth defects that can include heterotaxy and CHD^5,6,7^. While these findings suggest a role for metabolic regulation of LR patterning, as yet this remains poorly understood. Studies in mice showed hyperglycemia can impair left-right axis formation and heart tube looping, one of the first overt signs of visceral organ laterality^8^. This was associated with the disturbance of Nodal signaling, indicating glucose metabolism may be intimately involved in the specification of laterality.

Nodal and several other TGFβ family members, including *Lefty1* and *2,* are known to play critical roles in LR patterning^9,10,11,12^. In the mouse embryo, LR asymmetry is first noted by elevated expression of *Nodal* in crown cells at the left margin of the embryonic node, a transient bowl-shaped structure at the anterior tip of the primitive streak^13^. Cells in the node pit are mono-ciliated with motile cilia that generate right-to-left fluid flow. Cells at the node edge have primary cilia that senses flow, triggering calcium wave propagated to the embryo’s left side ^14,15,16,17,18,19,20^. This results in *Nodal* activation on the left, and *Lefty1* expression along the left-side of the floorplate, and *Lefty2* expression in the lateral plate mesoderm (LPM) on the embryo’s left-side^21, 22^. Subsequent left-sided expression of *Pitx2,* a homeobox containing transcription factor, provides visceral organ situs specification. In contrast, *Nodal* expression is repressed on the embryo’s right side by right-sided BMP signaling. This is accompanied by expression of another homeobox gene in the right LPM (R-LPM) – *Prrx1/prrx1a* in chick and zebrafish, respectively*. Prrx1* deficiency in chick or zebrafish, causes laterality defects^23^. *Snai1* is proposed to serve similar functions in mice, with *Snai1* KO mice showing randomization of heart looping^23, 24^.

To delineate the cell signaling pathways regulating LR patterning of organ situs during mammalian development, we conducted RNAseq on bisected mouse embryos at somite stages 3, 4, 5, and 6, spanning the time of *Nodal, Lefty2,* and *Pitx2* expression and just prior to heart tube formation and subsequent looping, one of the first sign of organ laterality. This is the first spatial transcriptome profiling of compartments of LR patterning, and data obtained has yielded novel insights into the molecular basis for patterning of visceral organ laterality.

## Results

To generate a spatial transcriptome profile of the compartments of LR patterning, mouse embryos at the 3, 4, 5, and 6 somite stages were microdissected and the embryonic node removed, followed by bisection of the embryo to generate right and left embryo halves for RNAseq analysis (Fig. 1a, b). This spans the developmental window for dynamic regulation of LR patterning, setting the stage for the first overt sign of visceral organ lateralization with heart looping at the 7 to 8 somite stages. Three embryos were collected at each somite stage, yielding RNAseq data for 24 half-embryo samples, 12 left and 12 right (Fig. 1a). To check the fidelity of the dissection, the RNAseq data were interrogated for *Nodal* transcripts, and as expected transcripts were observed only in the left half-embryos at the 3-5 somite stages (Fig. 1c). This was further verified by real time PCR quantitation of the RNA used for the RNAseq analysis (Fig. 1c). Similarly, strong left-sided expression of *Lefty2* transcripts was confirmed (Supplementary Figure 1). *Lefty1,* which is expressed only in a small population of cells on the left side of the floor plate, also showed left-sided expression. While this was not significant, likely due to its low abundance, further real time PCR analyses confirmed its left-sided expression (Supplementary Figure 1).

**Figure 1.**
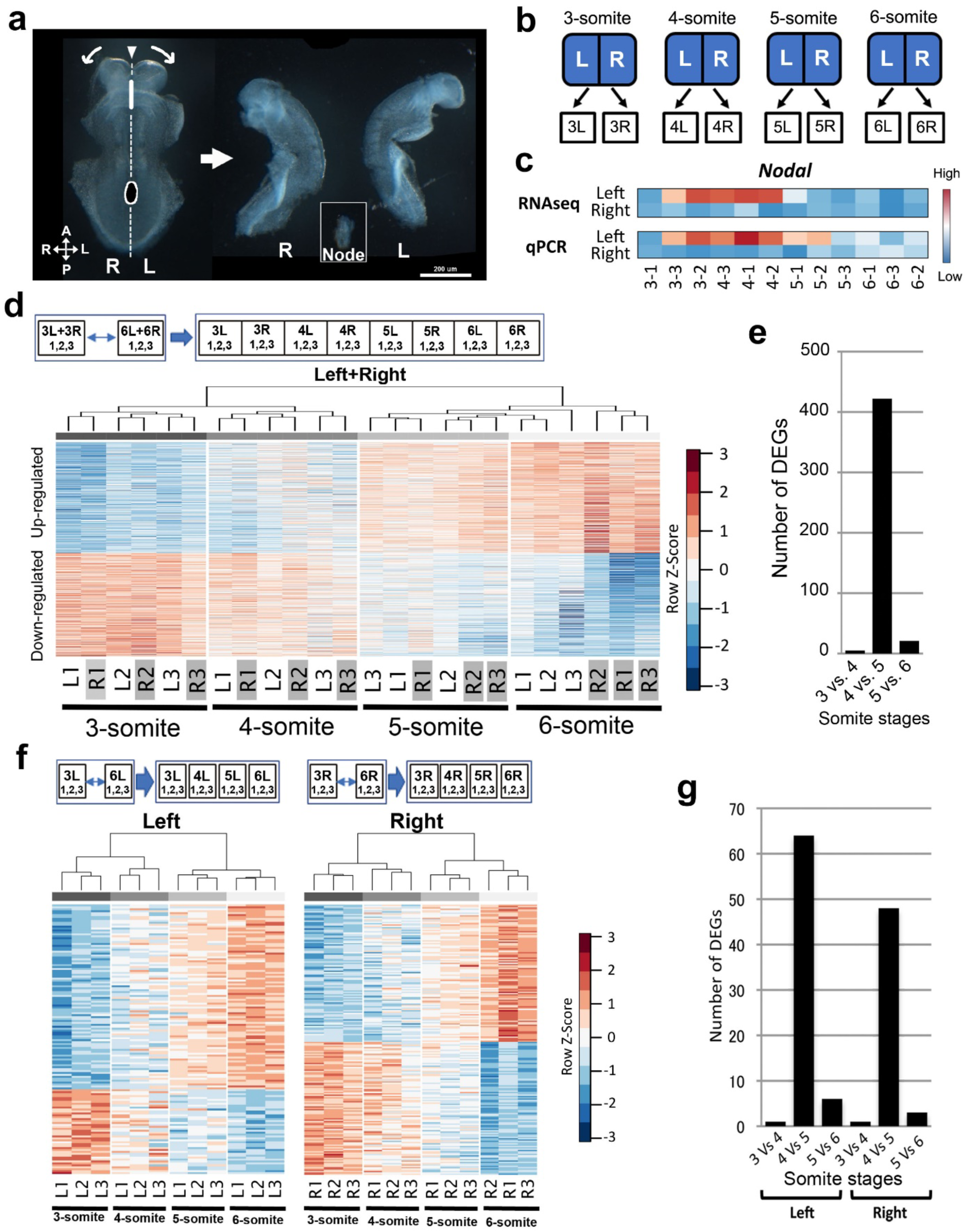
Transcriptome profile of differentially expressed genes at the 3-6 somite stages. (a) Embryos at the 3, 4, 5 and 6 somite stages were micro-dissected by first removing the embryonic node, followed by splitting the remaining embryo into left and right halves. (b) With three embryos harvested at each stage, this generated 24 half-embryo samples for RNAseq analyses, three left and three right at each somite stage, (c) Fidelity of dissection was verified with qPCR analysis for *Nodal* expression. (d) RNAseq data from the left and right half-embryos from the 3 and 6 somite stages were combined and differential gene expression analysis yielded 961 DEGs (1.5-fold up or down regulated; FDR <0.10). Self-organizing heat map of the 961 DEGs showed the samples clustered by embryo of origin at the 3 and 4 somite stages, but by the 6 somite stage, the samples clustered by L vs. R sides, indicating molecular diversification for left vs. right. (e) The number of DEGs between consecutive developmental stages is shown, revealing a peak between the 4 to 5 somite stage. (f) Diagram show analysis scheme used to recover left vs. right-sided DEGs. Left and right-sided DEGs between the 3 vs. 6 somite stage were recovered, and their expression at the 3, 4, 5, and 6 somite stages are shown in the heatmap. (g) Plot showing the number of left or right-sided DEGs observed between somite stages. The greatest change was observed between the 4 to 5 somite stage. This was observed for both the right and left half-embryo samples.

A bird’s eye view of the dynamic global transcriptional changes occurring during the time of LR visceral organ specification was obtained with construction of a heat map of gene expression changes (>1.5X fold, FDR≤1%) observed between the 3 vs. 6 somite stages in the left and right half-embryos combined (Fig. 1d). This yielded 961 differentially expressed genes (DEG), half up (n=497) and half (n=464) down-regulated (Fig. 1d; Supplementary Dataset S1). Hierarchical clustering of the DEGs from these 24 half-embryo samples automatically grouped them by somite stages (Fig. 1d). At the 3 and 4 somite stages, the half-embryos did not segregate by sides, but by embryo of origin. However, but by the 6-somite stage, the right vs. left half-embryos grouped together baed on whether it is from the right or left side, indicating molecular diversification distinguishing the embryo’s left vs. right sides (Fig. 1d). In comparison, at the 5-somite stage, the embryos appear to be in transition, with embryo 3 showing L vs. R segregation, but embryos 1 and 2 continuing to cluster by embryo of origin and not by left vs. right-sides. Overall, the largest transcriptional change occurred between the 4 to 5 somite stages with over 400 DEGs observed (Fig. 1e). DAVID Gene Ontology^25^. and Revigo analysis of these DEGs showed significant enrichment (FDR≤0.1) for genes associated with organogenesis (Supplementary Figure 2; Supplementary Dataset S2).

### Transcriptional Profiling the Left and Right Half-Embryos

To investigate the transcriptional profile of genes and pathways regulating development of the right or left half-embryos, DEGs for the left and right half-embryos at the 3 and 6 somite stages were recovered (Fig. 1f), yielding 140 DEGs for the left half-embryos (L-DEGs), and 206 DEGs for the right half-embryos (R-DEGs) (Supplementary Dataset S3). These 346 DEGs correctly sorted all 24 half-embryos by their somite stages (Figure 1f), and again the greatest number of DEGs was observed between the 4 to 5 somite stage. This was observed for both the left and right half-embryos (Fig. 1g). Enrichment analysis of up-regulated DEGs using DAVID Gene Ontology recovered the same five major pathways in both the left and right half-embryos (Supplementary Figure 3). This included heart development, skeletal system development, and tissue and embryonic morphogenesis (Supplementary Dataset S4). In contrast, pathways regulating anterior/posterior patterning and segmentation, and gland and blood vessel development were observed only on the right, and ossification and bone development were observed on the left, suggesting possible LR asymmetry related to these developmental processes (Supplementary Figure 3; Supplementary Dataset S4).

### Spatial Regulation of Nodal/Lefty and Bmp Signaling

To identify LR asymmetrically expressed genes, referred to as L vs. R-DEGs (L-DEGs/R-DEGs), we recovered transcripts that are differentially expressed in the left vs. right half-embryos at each somite stage (Fig. 2a). This yielded a total of 247 LR-DEGs across the four somite stages (Supplementary Dataset S5). Among the LR-DEGs recovered were many left-sided genes mediating TGFβ/BMP signaling, and as expected, this included *Nodal* and *Lefty2*, all genes highly expressed in the left half-embryo (Fig. 2b).

**Figure 2.**
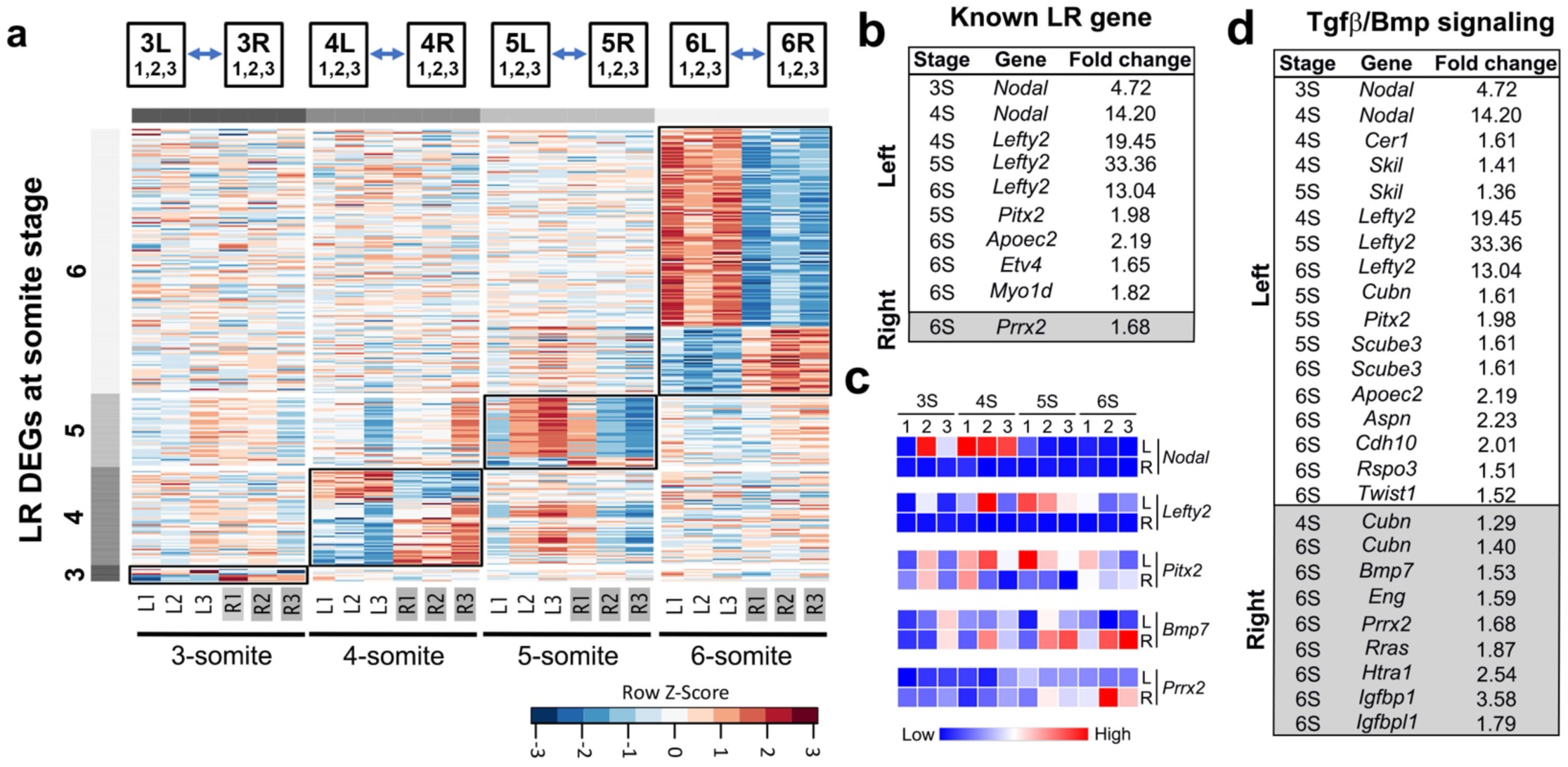
Left-right differential gene expression and genes associated with LR patterning. (a) Heatmap of DEGs between the left vs. right half-embryos. The samples included in the analysis are shown in the diagram above the heat map. (b) LR-DEGs known to play a role in left-right patterning. (c) Heatmap showing *Nodal, Lefty2, Pitx2, Bmp7,* and *Apobec2* transcript levels in the RNAseq data of the left vs. right half-embryos. (d) LR-DEGs associated with genes associated with TGFβ/BMP signaling, including genes known (*Nodal, Lefty2*), and not known to regulate LR patterning.

Also showing higher left-sided expression were other genes previously shown to regulate LR patterning including *Pitx2*, *Apobec2*^26^, *Etv4*^27^, and *Myo1d*^28, 29^.(Fig. 2b, c). These L-DEGs exhibited more modest 1.5 to 3-fold left-sided expression, and not the 4 to 30-fold higher left-sided expression observed for *Nodal* and *Lefty2* (Fig. 2b, c, and 2d; Supplementary Dataset S5). *Myo1d*, a non-muscle myosin, has been shown to regulate gut looping in *Drosophila*^28, 29^ and heart looping in zebrafish^30, 31^ and *Xenopus*^32^, but its LR asymmetric expression had not been reported previously. Left-sided expression was also observed for other genes in the TGFβ pathway not previously known to regulate LR patterning, such as *Skil*, *Twist1*, *Scube3*, and *Aspn* (Fig. 2d).

Also identified among the left-sided LR-DEGs was *Cer1*, a member of the Cerberus/DAN family of Tgfβ antagonists. Other members of this family have been shown to play a role in left-sided suppression of BMP signaling in the LPM^33^. Additionally, among genes mediating TGFβ/BMP signaling, were also several right-sided DEGs including *Bmp7* and *Prrx2,* and two other BMP pathway related genes, *Eng* and *Rras* (Fig. 2d). This is consistent with the known suppression of *Nodal* and activation of *prrx1a* expression by right-sided BMP signaling^23^. We note deficiency of *Prrx1* or *prrx1a* in chick and zebrafish respectively, has been shown to cause heart looping defects^23^. Also recovered as right-sided was expression of *Igfbp1* and *Igfbpl1*, insulin growth factor-like (IGF) binding proteins known to regulate IGF bioavailability and turnover. This suggests the possible modulation of IGF signaling, a pathway also likely subject to regulation by BMP7^34, 35^.

### Temporospatial regulation of glucose metabolism and IGF signaling

The right-sided expression of *Bmp7, Igfbp1* and *Igfbpl1,* would indicate a role for metabolism in the regulation of LR patterning. This likely involves modulation of glucose metabolism, as BMP7 is known to regulate glucose uptake and insulin sensitivity^34, 36^. A role for glycolysis in the regulation of LR patterning is supported by pathway enrichment analysis of the entire 24 half-embryo dataset, which yielded many glucose metabolism-related terms (Fig. 3a). These pathways were recovered largely from DEGs showing temporal regulation.

**Figure 3.**
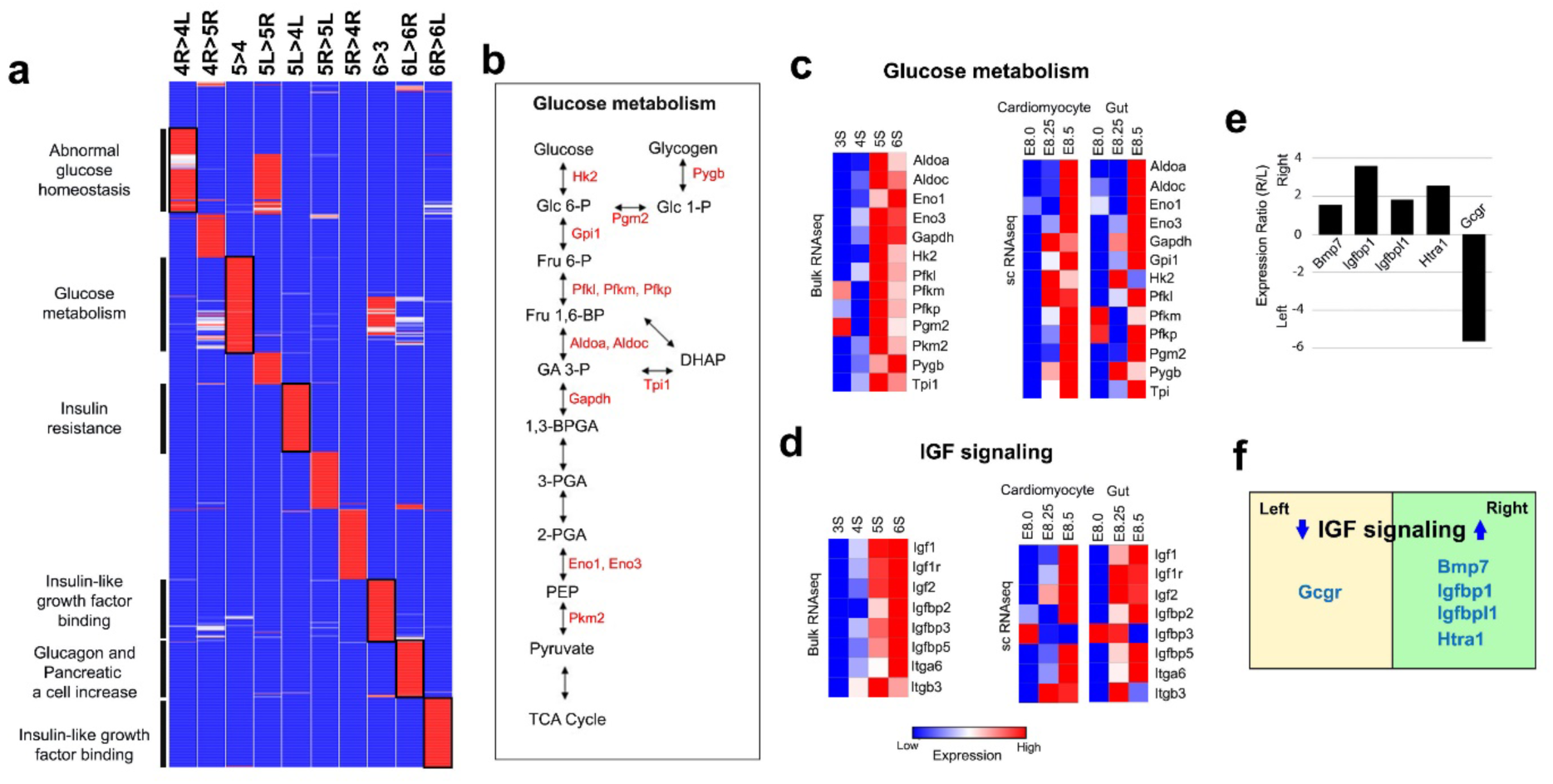
IGF signaling and Glucose metabolism. (a) ToppCluster analysis^94^ of DEGs recovered from comparisons between developmental stages or between left vs. right (indicated by column headers) organized as a heatmap generated using Morpheus (Broad Institute). DEGs that are higher (>) or lower (<) in expression between sides or between stages are as indicated. Notable is the recovery of many pathways related to IGF signaling and metabolism mostly involving comparisons between developmental stages. In contrast, cardiac related terms were only recovered in L vs. R comparisons, indicating LR asymmetry in heart development. The expected recovery of TGFβ signaling that is left sided was also observed. (b) Glycolysis pathway mediating glucose metabolism is shown, with upregulated glycolysis genes observed in the bulk RNAseq shown in red. (c,d) Heatmap of transcripts observed in the bulk/sc RNAseq analysis related to genes regulating glucose metabolism (c), and IGF signaling (d). scRNAseq data in cardiomyocytes and the gut progenitors are from previously published mouse embryo single cell RNAseq data^40^. (e, f) LR-DEGs recovered related to IGF signaling, showing their fold change (g) and left vs. right-sided distribution (h).

Almost all the genes mediating glycolysis were recovered, including phosphofructose kinase (PFK), the key allosteric enzyme regulating glycolysis (Fig.3b). All three PFK isoforms are highly expressed (*Pfkp*, *Pfkl*, *Pfkm*; Fig. 3c). Pathway enrichment analysis yielded glucose metabolism (-Log10 p-value =10), abnormal glucose homeostasis (-Log10 p-value =2.6), insulin resistance (-Log10 p-value =2.15), insulin-like growth factor (IGF) binding (-Log10 p-value =4.24), increased glucagon (-Log10 p-value =3.27), increase pancreatic a cells producing glucagon (-Log10 p-value =3.1), and other metabolic pathways (Fig. 3a, Supplementary Figure 4). All the DEGs regulating glucose metabolism and most of those mediating IGF signaling showed elevated expression starting at the 5-somite stage (Fig. 3c,d), coinciding with genome-wide transcriptional reprogramming observed at the 4 to 5 somite stage (Fig. 1d-g).

In addition to right sided expression of *Bmp7, Igfbp1* and *Igfbpl1,* we also observed right-sided expression of *Htra1,* a gene that regulates the availability of insulin-like growth factors via binding and cleavage of insulin-like growth factor binding proteins (Fig. 2c, 3e,f)^37^. As Htra1 can also bind and inhibit TGFβ family members^38, 39^, this suggests a role in mediating cross talk between BMP and IGF signaling and the right-sided suppression of TGFβ signaling. Interestingly, left-sided expression was observed for the glucagon receptor and other genes associated with pancreatic a cell increase, suggesting possible LR asymmetry in development of the pancreas (Fig. 3a,e,f)Together these findings support the metabolic regulation of glycolysis playing a role in LR patterning of visceral organ situs.

### Metabolic regulation in the cardiomyocyte and gut lineages

Looping of the embryonic heart and gut tubes are notable indicators of visceral organ situs specification. Hence, we further investigated the expression of DEGs involved in glucose metabolism and IGF signaling in differentiating cardiomyocyte and gut lineages using publicly available scRNAseq dataset derived from mouse embryos at E8.0 to 8.5, estimated to be at the 0-7 somite stages^40^. The cardiomyocytes and gut forming cells both showed abundant expression of genes mediating glycolysis and IGF signaling at E8.25 and E8.5, but not at E8.0 (Fig. 3c,d; Supplementary Figure 4), consistent with our bulk RNAseq analysis showing a global metabolic transition at the 5-somite stage (Fig. 1c). It is notable that nearly all the genes mediating glycolysis were recovered in both the bulk and single cell RNAseq data (Fig.3c). All the DEGs regulating glucose metabolism and most of those mediating IGF signaling showed elevated expression from the 5-somite stage (Fig. 3c,d), coinciding with the time of genome-wide transcriptional reprogramming (Fig. 1d-g). These findings indicate genes regulating glycolysis are temporally regulated in both the cardiomyocyte and gut lineages.

### LR-DEGs and Heart Development

Development of the heart involves migration of cardiac progenitors derived from the right and left lateral plate mesoderm (R-LPM/L-LPM) to the midline, followed by their coalescence to form the primitive heart tube. Its subsequent right-sided looping at the 7 to 8 somite stage provides one of the first overt sign of visceral organ laterality. Examination of LR-DEGs yielded many genes related to heart development, with most being left-sided, suggesting left bias in cardiomyocyte differentiation (Fig. 4a; Supplementary Dataset S1, S2) This included the transcription factor *Mef2c*, ventricular sarcomeric proteins such as myosin heavy chain 7 (*Myh7*) and cardiac troponin c (*Tnnc1*), and ryanodine receptor 2 (*Ryr2*), a calcium channel protein mediating excitation-contraction coupling. (Fig.4a). A few right-sided DEGs associated with heart development were also identified, including *Gja5*, a gap junction protein expressed in endothelial cells and also the cardiac conduction system (Fig. 4a). This is consistent with previous studies showing cardiac pacemaker cells arise from the R-LPM^41^. Also, right-sided was expression of e*ndoglin* (*Eng*), *neuregulin* (*Nrg1*)*, heparan sulfate proteoglycan2* (*Hspg2), g*enes known to regulate cardiac cushion mesenchyme development (Fig. 4a). The right-sided expression of *Prrx2*, a gene known to regulate epithelial-mesenchymal transition (EMT), also may play a role in endocardial EMT and development of cardiac valves. These right-sided DEGs together with *Rras* would suggest enhancement of blood vessel development on the right (Fig.4a), consistent with recovery of blood vessel development from the R-DEG pathway enrichment analysis (-Log10 p-value<6.79; Figure S3).

**Figure 4.**
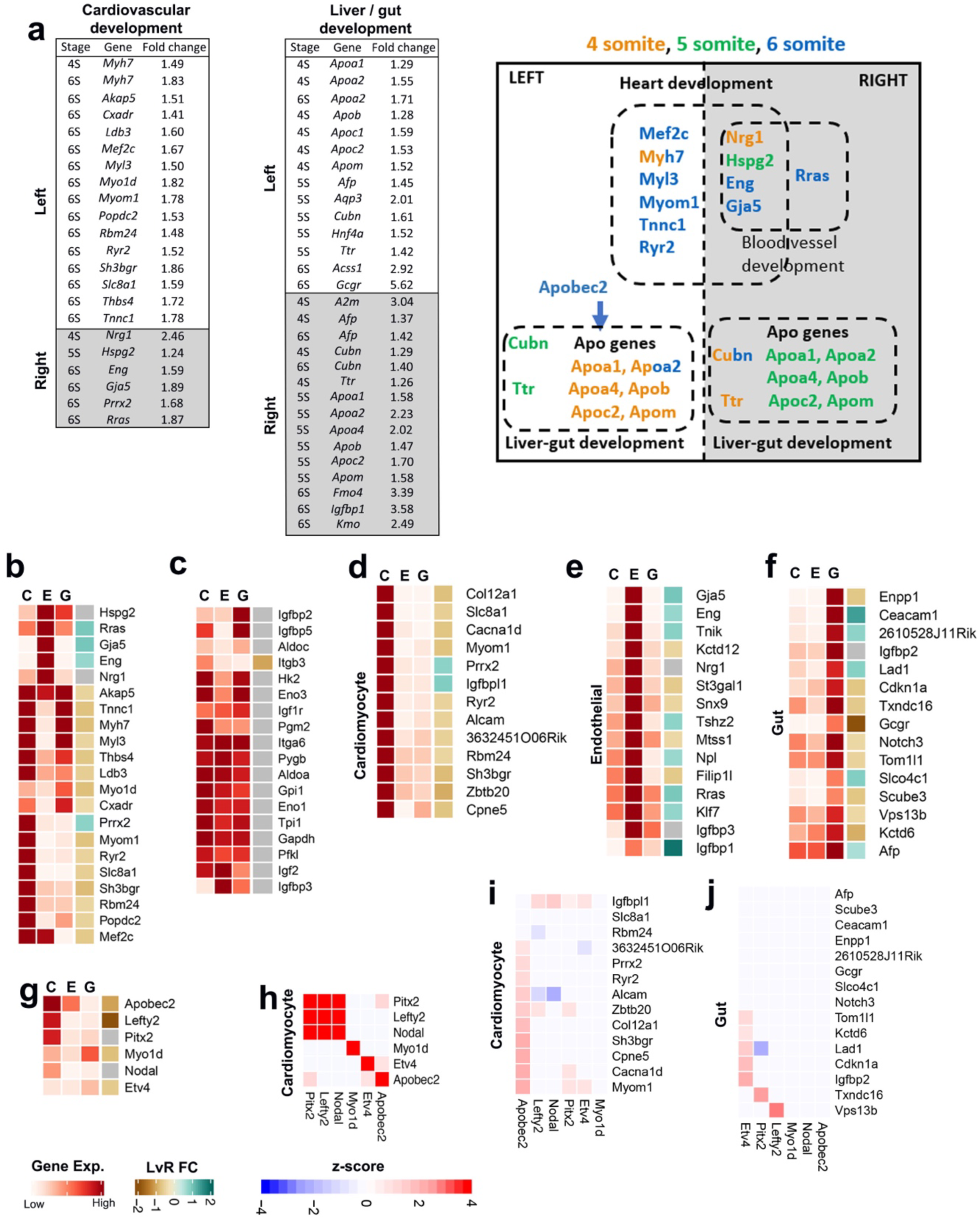
Expression of LR-DEGs in heart and gut development. (a) Tables summarizes expression of LR-DEGs related to heart and liver/gut development, and their L vs. R sided and somite stage of expression. This is further illustrated for selected heart or liver/gut developmental genes in the diagram on the right, with by font color indicating somite stage of expression. (b-j) Analysis of the expression of DEGs identified in the bulk RNAseq in the cardiomyocyte, endothelial and gut cell lineages using previously published single cell RNAseq data^40^. (b). Heat map of DEGs for cardiovascular development from panel (a) as observed in the cardiomyocyte, gut and endothelial cell lineages using previously published scRNASeq data^40^. (c). Heat map of DEGs associated with IGF signaling and glucose metabolism in the cardiomyocyte, gut and endothelial cell lineages using previously published scRNAseq data^40^. (d-f) Heat map showing expression of DEGs identified as cardiomyocyte (d), endothelial (e), or gut (f) specific in the cardiomyocyte, endothelial, and gut cell lineages using previously published scRNAseq data^40^. (g) Expression of genes regulating LR patterning in the cardiomyocyte, endothelial and gut cell lineages using previously published scRNAseq data^40^. (h) Co-expression of known LR genes in the cardiomyocyte cell lineages^40^. (i,j) Co-expression of known LR genes with genes specific to the cardiomyocyte (i) or gut/endodermal (j) lineages using previously published scRNAseq data^40^.

### LR DEGs Associated with Liver and Gut

The endoderm gives rise to the gut tube at E8.75, and from which other organs emerge such as the liver and pancreas. Several notable liver/gut related genes were recovered among the LR-DEGs, including multiple apolipoproteins (*Apoa1*, *Apoa2*, *Apoa4*, *Apob*, *Apoc2*, *Apom*), *Cubilin* (*Cubn*) - an endocytic receptor for high-density lipoproteins and other ligands, and transthyretin (*Ttr*) - a protein mediating transport of thyroid hormone, retinol, and other ligands (Fig. 4a). The expression of these genes shows temporospatial regulation, with the apolipoprotein genes exhibiting right-sided expression at the 4-somite stage, and left-sided expression at the 6-somite stage (Fig. 4a). Interestingly, higher left-sided expression was also observed for apolipoprotein B mRNA editing enzyme catalytic polypeptide 2, *Apobec2*, a putative cytidine deaminase previously shown to modulate TGFβ signaling in LR patterning^26^.

### LR-DEGs Expression in Cardiomyocytes, Endothelial and Gut Cell Lineages

LR-DEGs associated with cardiovascular development were further examined for their expression in different cell lineages using the same publicly available scRNAseq dataset described above^40^. In cardiomyocytes, high-level expression was observed for nearly all the L-DEGs, and for the R-DEGs, only *Prrx2* was highly expressed (Fig. 4b). The remaining five R-DEGs, *Hspg2, Rras, Gja5, Eng, Nrg1, s*howed strong expression in the endothelial lineage, consistent with a role in blood vessel development and in development of the cardiac endocardial cushions (Fig. 4b). Interestingly, the gut cell lineage also showed strong expression of some of the cardiomyocyte associated L-DEGs, including contractile proteins such as *Myh7, Myl3, Tnnc1, Myo1d* (Fig. 4b). In contrast to the restriction of many LR-DEGs to specific cell types, expression of glycolysis-related metabolic genes showed moderate to high level expression in all three lineages (Fig. 4c), consistent with the global activation observed in the spatial transcriptomic analyses (Fig. 3a). It is worth noting among the highly expressed metabolic genes are *Igf2* and *Igf1r*, ligand and receptor mediating IGF signaling previously shown to be essential for growth of the embryo^42^, and the embryonic heart^43,44,45^.

To further investigate LR DEGs that might have lineage specific roles, we interrogated the scRNAseq data for LR-DEGs highly expressed only in the cardiomyocyte, endothelial, or gut forming cells (Fig. 4d-f; Supplementary Figure 4). In the cardiomyocytes, all but two of the cardiomyocyte-specific LR DEGs were L-DEG (Fig.4d), supporting the left-bias in heart development observed by the bulk RNAseq analysis. There also two R-DEG, *Prrx2* and *Igfbpl1*, among the cardiomyocyte specific DEGs (Fig. 4d). In contrast to the predominance of L-DEGs in the cardiomyocytes, the endothelial or gut-specific LR-DEGs included many R and L-DEGs (Fig. 4e, f). Among genes with a role in the regulation of IGF signaling, the R-DEG *Igfbp1* was observed in the endothelial lineage (Fig. 4e), while the L-DEG *Gcgr* encoding the glucagon receptor was observed in the gut (Fig.4f).

Analysis for expression of the LR patterning genes in the cardiomyocyte, endothelial and gut lineages showed *Apobec2, Lefty2,* and *Pitx2* are strongly expressed in the cardiomyocytes, while *Apobec2* is observed in the endothelial cells, and *Myo1d* in the gut (Fig. 4g). These findings support *Lefty* and *Pitx2* having a predominant role in patterning cardiac laterality. Consistent with this, *Nodal, Lefty2* and *Pitx2* were found to be coexpressed in the cardiomyocytes, but this was not observed for the other LR patterning genes (*Etv4, Myo1d*, *Apobec2*) (Fig. 4h). However, *Nodal*, *Lefty2*, *Pitx2* were not coexpressed with the cardiomyocyte specific LR-DEGs, such as *Myh7*, *Myom1*, *Ryr2*, *Nkx2-5*, *etc* (Fig. 4i, data not shown). Instead, the latter LR-DEGs showed low level co-expression with *Apobec2* (Fig. 4i). In the gut lineage, the gut-specific LR-DEGs showed low level coexpression with *Etv4*, but none were coexpressed with *Myo1d* (Fig. 4j). Additionally, *Vps13b* showed coexpression with *Lefty2*, and *Txndc16* with *Pitx2* (Fig.4j).

### *Myo1d* Regulation of Gut Looping

The novel observation of *Myo1d* as a left-sided DEG and the finding of its specific expression in the gut lineage in the mouse embryo prompted us to further investigate its role in LR patterning in mice and man. Using *in situ* hybridization analysis, we confirmed transient left-sided expression of *Myo1d* transcripts in the early postimplantation mouse embryo (Fig. 5a). *In vitro* lentiviral mediated shRNA knockdown of *Myo1d* in cultured mouse embryos caused abnormal heart looping, indicating evolutionary conservation of a role for *Myo1d* in regulating laterality in the mouse as has been observed in zebrafish, *Xenopus*, and *Drosophila* mutants^28,29,30,31,32^(Fig. 5b). To further investigate the role of *Myo1d* in LR patterning, we generated *Myo1d* knockout (KO) mice using CRISPR gene editing. The *Myo1d* KO mice are homozygous viable, and while they showed no overt heart or lung situs defects, reversal of gut looping was observed with analysis of sigmoid colon orientation relative to the superior mesenteric artery (Fig. 5c; p<0.01). We also examined gut looping in *myo1d* knockout zebrafish. As *myo1d* is expressed as maternal transcripts, maternal zygotic (MZ) mutants were generated to produce true *myo1d* null zebrafish embryos. Abnormal gut looping was observed in these *myo1d* deficient zebrafish mutants comprising of right loop, straight, or misaligned gut looping phenotypes (Fig. 5d, e). While all the zebrafish mutant embryos were initially viable, those with abnormal gut looping died between 7 to 14 days post fertilization with an odds ratio of 2.5-2.6 (p<0.0001). This suggests the abnormal gut looping is associated with gut obstruction that is incompatible with long term viability (Fig. 5f).

**Figure 5.**
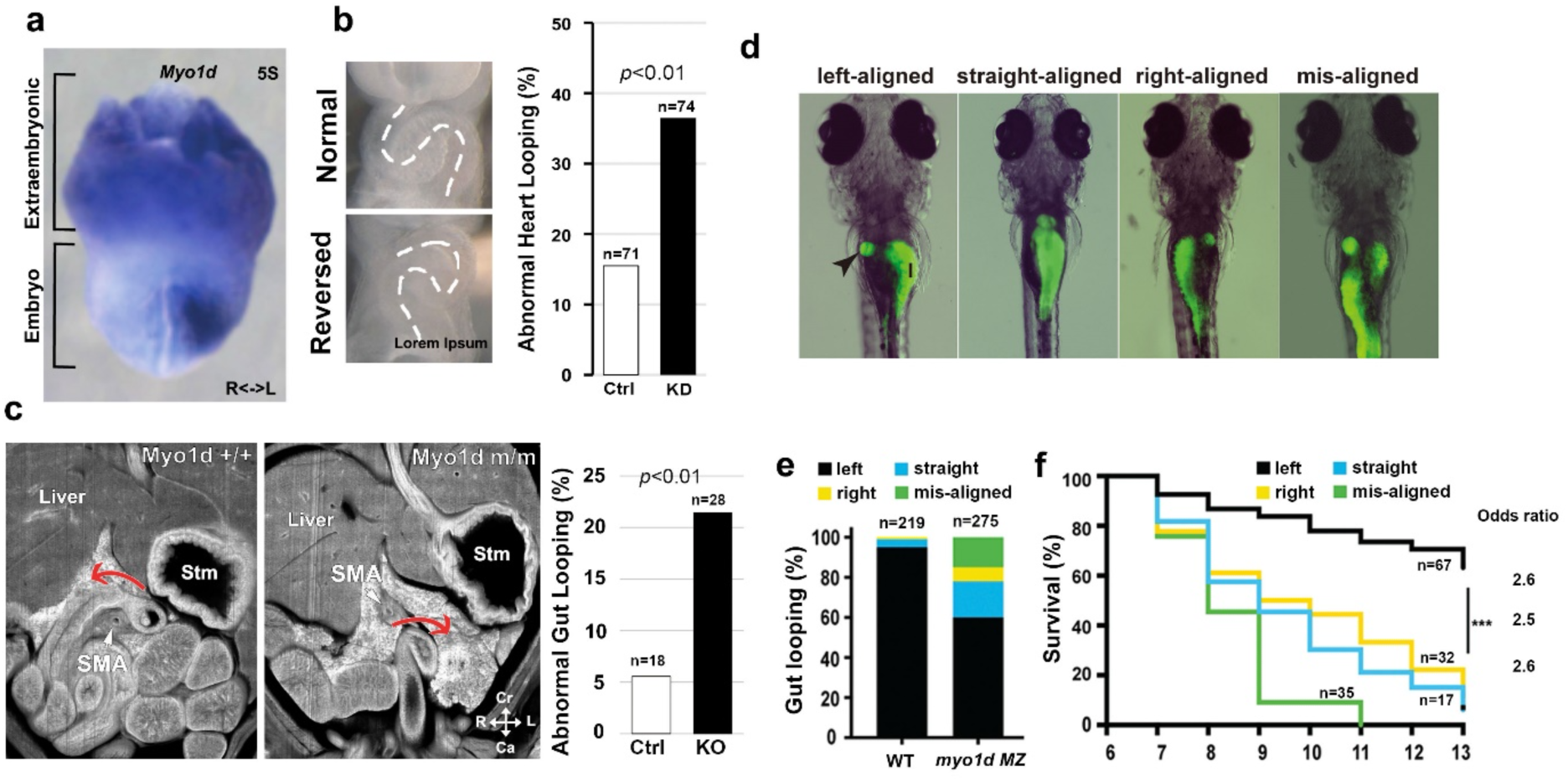
*Myo1d* deficiency causes gut looping defects in mice and zebrafish. (a) *In situ* hybridization showing left-sided expression of *Myo1d* in the early mouse embryo. (b) Lentiviral mediated *Myo1d* siRNA knockdown in mouse embryos cultured in vitro caused significant increase in reversed/abnormal heart looping. (c) Analysis of the sigmoid colon looping orientation relative to the superior mesenteric artery (SMA) of *Myo1d* KO mice showed significant gut looping defect. Rather than normal counterclockwise looping, the KO embryos showed significant increase in abnormal clockwise looping. (d) Examination of gut looping in zebrafish larvae at 6dpf after feeding with PED6. Visualization of the gut revealed various defects in gut looping. Arrowhead denotes gallbladder. I: Intestine. (e) *myo1d* maternal zygotic (MZ) larvae showed increased incidence of abnormal gut looping compared to wildtype. I = Intestine; arrowhead denotes gallbladder. (f) Survival analysis showed marked loss of *myo1d MZ* larvae with abnormal gut looping. ***p<0.0001 with Chi square analysis (gut looping phenotype distribution: n=67 left, 32 straight, 17 right, 35 mis-aligned).

### *MYO1D* Mutation Clinically Associated with Heterotaxy and Gut Malrotation

To investigate the clinical relevance of *MYO1D* mutations in gut looping abnormalities, we pursued whole exome sequencing analysis of 73 heterotaxy patients with CHD. Heterotaxy in this patient cohort is defined as abnormal situs involving the cardiovascular system with or without lung or abdominal situs abnormalities^46^. Two heterotaxy patients with CHD (7007; 7101) were each observed to have two rare *MYO1D* missense mutations. Genotyping of parents showed the two mutations in each subject are in cis, thus indicating both patients are heterozygous for a single mutant *MYO1D* allele comprising two linked missense mutations (Fig. 6a; Supplementary Table S1). Patient 7101 has normal atria and abdominal situs, but with an inverted L-loop ventricle associated with a double inlet left ventricle. Patient 7007 at 21 years of age had emergency surgery for intestinal malrotation associated with internal herniation of the omentum (Fig. 6b-d). This patient also had CHD comprising left atrial isomerism, interrupted inferior vena cava, and unbalanced atrioventricular septal defect (Fig. 6e). Organ laterality defects observed included a transverse liver, intestinal malrotation with the entire small bowel on the right and colon on the left side of the abdomen (Fig. 6d), and polysplenia (Fig. 6f). These observations support a conserved role for *MYO1D* in gut looping from *Drosophila* to zebrafish, mouse, and man^28,29,30,31,32^.

**Figure. 6.**
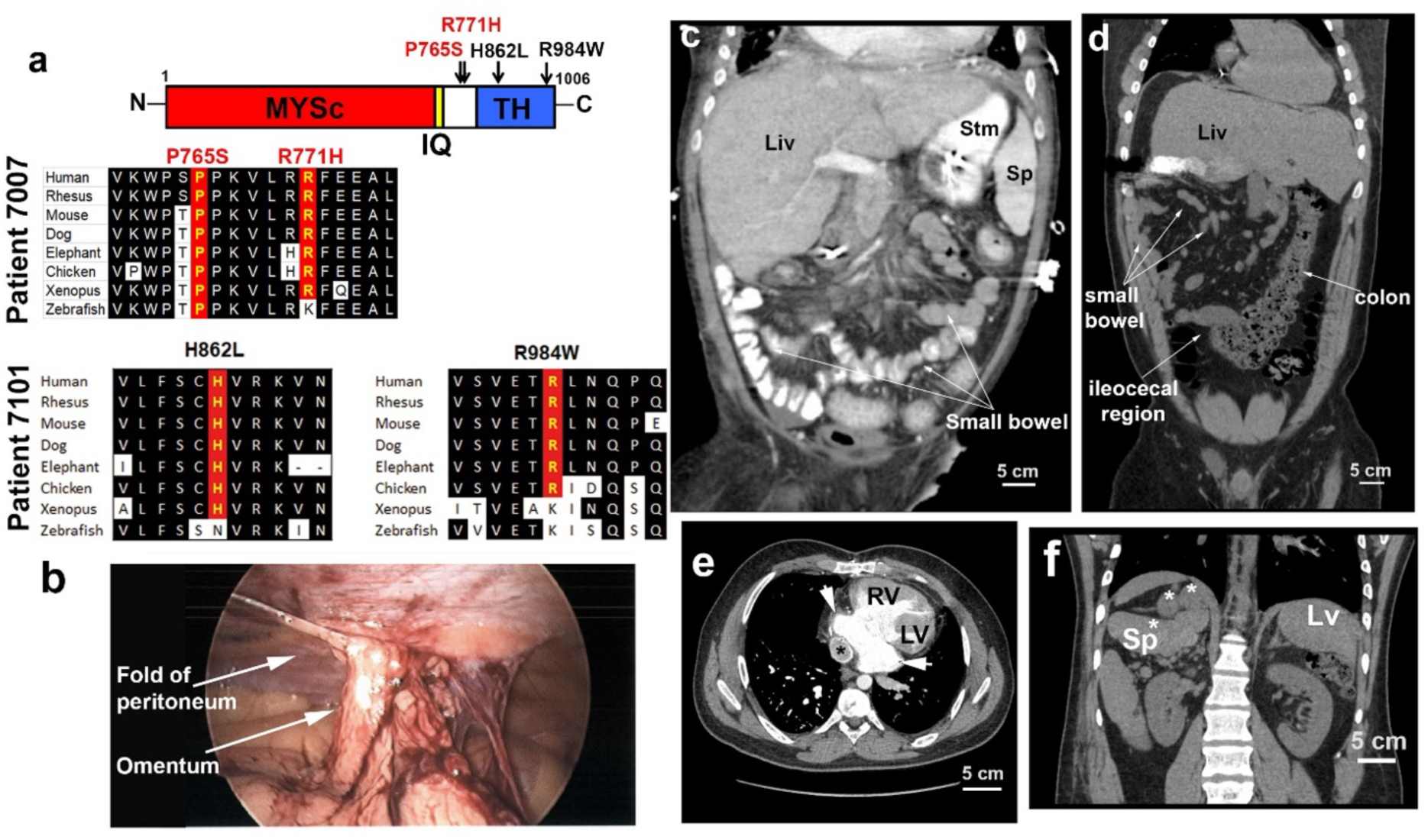
*MYO1D* deficiency causes heterotaxy in human. (a) Two *MYO1D* mutations in cis were found in each of two heterotaxy patients (top). Amino acid sequence alignment of the mutations in patient 7007 with gut malrotation showed high sequence conservation. (b) Laparoscopy of patient 7007 demonstrated malrotation with internal hernia. The abnormal peritoneum fold created a defect between itself and the anterior abdominal wall, with the omentum herniating into this defect. (c) Computed tomography (CT) of individual with normal gut rotation showed small bowel in center of abdomen and colon on the right. (d) Computed tomography of patient 7007 shows intestinal malrotation. The colon was entirely in the left abdomen, with ileocecal region in the center. Multiple small bowel loops were visible in the right abdomen. (e,f) Chest CT scan (e) of heterotaxy patient 7007 revealed unbalanced atrioventricular septal defect with dominant right ventricle (RV). Bilateral broad based left atrial appendages are observed (white arrow indicate left atrial isomerism). Polysplenia was observed in the right abdomen (f). * denotes Fontan conduit. LV, left ventricle; RV, right ventricle; Sp, spleen

## Discussion

We generated the first spatial transcriptome profile of LR patterning and recovered TGFβ signaling mediated by Nodal/Lefty as the primary determinant of the LR body axis. No other cell signaling pathway was detect with the 10-30 fold left-sided expression seen for *Nodal/Lefty2*. Also observed was expression of the homeobox transcription factors *Pitx2*, on the left and *Prrx2*, on the right. Both showed modest change in LR expression - 2 and 1.7 fold, respectively. We also noted right sided expression of *Bmp7,* which is known to regulate glycolysis and insulin sensitivity, and also transcripts for several IGF binding proteins (1.8 to 3.6) (Fig.7). These findings are consistent with prior studies indicating two parallel conserved pathways patterning organ laterality comprising left-sided expression of *Nodal/Lefty*/*Pitx2* and right-sided expression of *BMP/Prrx* (Supplementary Figure 6a). In chick and zebrafish embryos, previous studies showed left-right patterning is associated with Bmp signaling and transient right-sided *Prrx1/prrx1a* expression. *Prrx1/prrx1a* deficiency was shown to cause laterality disturbance with mesocardia^23^. While laterality defects are not observed *in Prrx1* and *Prrx1/Prrx2* KO mice^47, 48^, this may reflect functional redundancies, such as by *Snai1. Snai1* deficient mice exhibit laterality defects with randomized heart looping^24^. Both *Prrx1* and *Snai1* regulate epithelial-mesenchymal transition and cell migration, cellular processes with demonstrated roles in heart development and looping of the heart tube^23, 49^.

**Figure 7.**
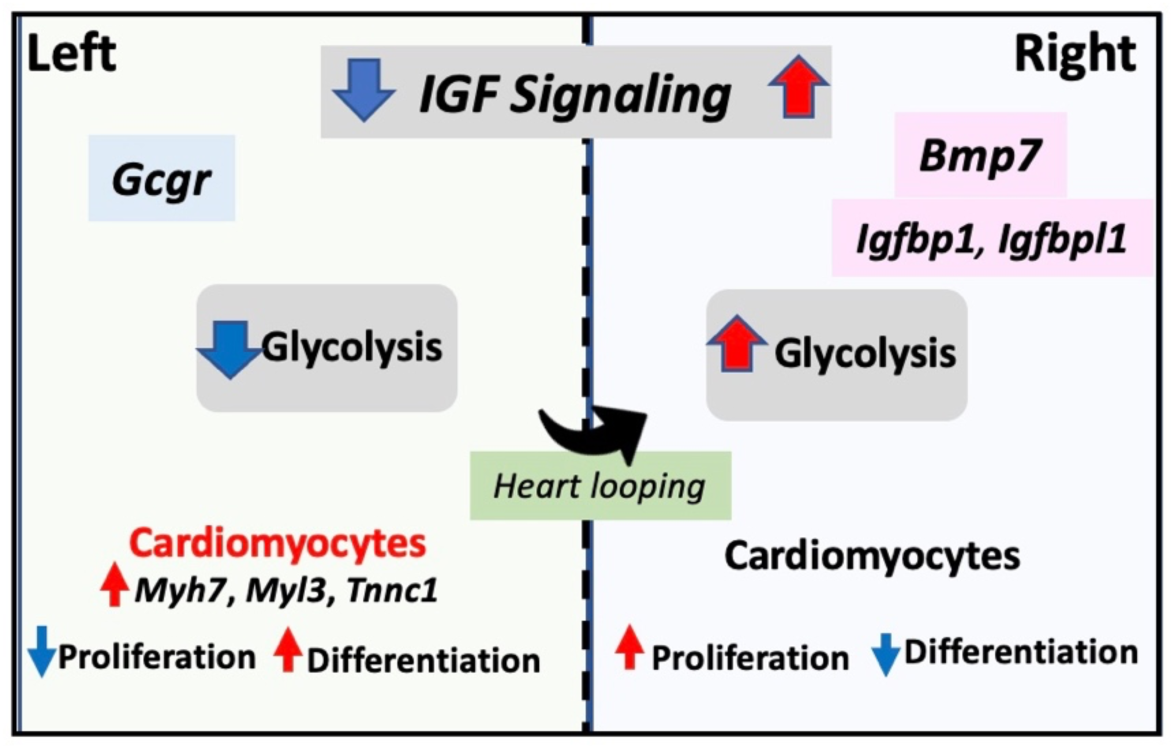
Metabolic regulation of LR patterning via IGF signaling. Right-sided bias in Bmp7 expression stimulates IGF signaling and glycolysis on the embryo’s right, promoting proliferation, but repressing differentiation. The reverse is expected on the left with less proliferation and more differentiation. This can account for many cardiomyocyte differentiation markers being left sided. This asymmetry in proliferation and differentiation may also contribute to LR patterning of liver/lung lobation

Coexpression of *Nodal/Lefty/Pitx2* was observed in the cardiomyocyte lineage. This is consistent with the LPM being both the site of *Nodal/Lefty2* expression and also the source of cardiac progenitors. The scRNAseq analysis indicated a role for Nodal/Lefty signaling in the early cardiomyocyte progenitors, while transient Apobec2 expression was observed in the differentiated cardiomyocytes. Apobec2 has been shown to be part of a HDAC transcriptional complex and thus may further transcriptionally regulate LR patterning of cardiomyocyte differentiation^50^. We also recovered, *Cer1*, a Cereberus-related BMP antagonist as a left-sided DEG at the 4 somite stage. As *Dand5 (Cerl2)* has been shown to mediate Nodal repression^51, 52^, perhaps *Cer1* expression may promote cessation of *Nodal* expression at the 5-somite stage. Several other TGFβ family members were also recovered as L-DEGs including *Skil*, *Twist1*, *Scube3*, *Aspn,* but their role in LR patterning has not been investigated.

Unexpectedly, we identified as a novel L-DEG, *Myo1d*, a gene known to regulate gut looping in *Drosophila*^28, 29^. Our analysis of scRNAseq data suggests while *Myo1d* may regulate early events in patterning of gut laterality, *Etv4* expression may underlie later LR patterning regulation of the differentiating gut lineage. Importantly, we showed *Myo1d* has a conserved role in regulating gut looping in mice, zebrafish and humans. As *Myo1d* expression was observed in the gut forming cells in the embryo, this would suggest this nonmuscle myosin may exert cell-autonomous function in gut looping. We observed *Myo1d* expression in cardiomyocytes, and while heart looping defect was noted after acute *Myo1d* knockdown with shRNA treatment, *Myo1d* knockout mice did not show heart looping defects, suggesting possible functional redundancies mediated by other myosins.

The most striking finding from our transcriptome profiling was evidence for the metabolic regulation of LR patterning. This is indicated by right-sided expression of *Bmp7, Igfbp1*, *Igfbpl1,* and *Htra1.* Studies in diabetic mice have shown *Bmp7* can augment insulin sensitivity with enhanced glucose uptake and increase in glycolysis^34^. As glycolysis can stimulate cardiomyocyte proliferation while suppressing differentiation^53^, this would predict enhanced cardiomyocyte differentiation on the left relative to the right consistent with finding of more left-sided cardiomyocyte differentiation in our transcriptome profiling (Fig. 7). LR asymmetry in proliferation, with higher proliferation in cardiac progenitors on the right has been observed in the early mouse embryo right^54,55,56^. The right-sided metabolic repression of cardiomyocyte differentiation together with *Prrx2* regulated EMT and cell migration may contribute to rightward looping of the heart tube (Fig. 7)^23^. As glycolysis also similarly regulates proliferation and differentiation of endodermal derivatives^57, 58^, this may help pattern liver/lung lobation, a process likely mediated by tissue growth regulation by IGF signaling^42^.

These findings indicating possible metabolic regulation of LR patterning provides novel mechanistic insight into the known association of pregestational diabetes (PGD) with birth defects involving the disturbance of LR patterning^59, 60^. This is well documented by a large birth cohort study comprising over 50,000 infants that reported OR of 12.3 (7.3-20.5) for heterotaxy with CHD^61^. Clinical^62, 63^ and animal studies ^64,65,66,67^ have provided evidence that hyperglycemia is the underlying cause for PGD embryopathy, consistent with our findings indicating the modulation of glycolysis in embryonic LR patterning. Notable in this regard is the previous identification of a deletion CNV encompassing *PFKP* as a genetic cause for heterotaxy^68^. *PFKP* encodes phosphofructose kinase, the key allosteric enzyme regulating glycolysis, and its morpholino knockdown in *Xenopus* embryos resulted in heterotaxy^68^. Additionally, *ZIC3*, a gene well described to cause heterotaxy, both clinically^69^ and in mice^70, 71^, was recently shown to regulate glycolysis required for the maintenance of plruipotency^72^, with *Zic3* deficiency causing reduction in glycolysis from failure to activate genes mediating glycolysis^72^. Significantly, the deletion of *Zic3* in the LPM disrupted LR patterning^73^, indicating the LPM is likely one of the target tissues under metabolic regulation for LR patterning. As the cardiac progenitors are derived from the LPM, this may explain the common finding of complex CHD with heterotaxy in patients with *Zic3* mutations. The role of metabolism in left-right patterning is further suggested by a recent study in which single cell mass spectrometry showed different metabolites are enriched in the left vs. right blastomeres of the 8-cell *Xenopus* embryo^74^.

Overall, these findings suggest dysregulated glucose metabolism may underlie the high incidence of birth defects associated with LR patterning disturbance in PGD. We estimate the sensitive developmental window for PGD embryopathy in human development at 3 weeks gestation or Carnegie stages 7/8, equivalent to the 4-6 somite stage mouse embryo^75^. This suggests achieving glycemic control to prevent PGD embryopathy would be difficult, given 50% of pregnancies are unplanned^76^. With the expected increase in PGD embryopathy with the continuing rise in type 2 diabetes^77^, the overall incidence of birth defects will continue to rise. As birth defects involving CHD associated with heterotaxy have disproportionately higher morbidity and mortality^2^, there also will be significant increase in clinical burden for infants born with PGD embryopathy. However, before new therapeutic strategies can be developed for prevention of PGD associated birth defects, more mechanistic insights are needed into how metabolism affects transcriptional programming of LR patterning and embryonic development. Overall, the unique transcriptome dataset generated in this study will be invaluable for further investigating the pathomechanisms of birth defects associated with laterality disturbance.

## Methods

### Animal Protocol Approval

All animal procedures were conducted according to relevant national and international guidelines (AALAC and IACUC) and have been approved by the University of Pittsburgh (Protocol mouse #15127136, Protocol zebrafish# 16088842) and Jackson Laboratory Animal Care and Use Committee (Protocol #99066).

### Embryo Microdissection

The embryonic node was first removed, followed by a cut along the midline at the posterior margin of the anterior intestinal portal, splitting the heart field (3 and 4-somite)/primitive heart tube (5-6 somite). This is followed by a cut at the anterior-midline of the head fold and then using forceps to gently hold the left and right sides the head fold and pulling in opposite direction, splitting the embryo along the midline into left and right halves. All experiments were carried out using protocols approved by the Institutional Animal Care and Use at the University of Pittsburgh.

### Transcript and RNAseq Analysis

Total RNA from mouse embryos were isolated using RNeasy plus micro kit with on-column DNase I digestion method (QIAGEN). cDNA libraries were produced by using an Ovation RNAseq amplification kit Ver. 2 (NuGen).

Real-time PCR was performed using Applied Biosystems HT7900 with Power SYBR Green PCR Master Mix (Life Technologies) following standard protocol. The ubiquitously expressed *β-actin* gene was used as an internal control.

For RNAseq, cDNA library was sequenced using SOLiD 5500xl (3-6 somite stage) (BGI Americas). Alignment and gene-level quantification was performed using the mouse reference genome mm9 (check) and the LifeScope Software suite for the 3-6 somite stage data and TopHat2 (v2.0.9)^78^ and HTSeq-count (v0.5.4p5)^79^ for the 1 somite stage data. We considered the intersection of significantly differentially expressed (DE) genes identified by DEseq (v1.12.0)^80^ and edgeR (v3.2.3)^81^ as follows.

For 3-6 somite data, dispersion parameters were estimated accounting for side (L or R) and embryonic stage (3, 4, 5 or 6) as factors. For edgeR, for each gene, the maximum of trended and tag-wise dispersion estimates was used. For DESeq, the “pooled-CR” method with the “maximum” sharing mode was used for estimating dispersions. Genes with a false discovery rate (Benjamini-Hochberg) <= 0.2 in both methods were considered significantly differentially expressed. Gene Ontology (GO) term enrichment analysis was performed using DAVID (v 6.7)^82^ and data were visualized by REVIGO^83^.

### Analysis of Single Cell RNA-sequencing Data

Single cell RNA sequencing data of E8.5 mice^84^ were downloaded from Array Express (https://www.ebi.ac.uk/arrayexpress/, accession: E-MTAB-6153). Normalized counts were read into the R programming language (https://www.r-project.org) and processed for further analysis. Cluster/cell-type annotations were taken from the original publication and we defined the expression prevalence of a gene–cluster–combination as the fraction of cells expression a gene in a cluster, divided by the overall fraction of cells expressing that gene. Expression prevalence was calculated using all 19,386 cells and 33 clusters for the genes shown in Supplementary Figure 4, where gene/cluster combinations with less than three gene-expressing cells were omitted (grayed out). Heatmaps were generated with the heatmap function from the pheatmap package (https://CRAN.R-project.org/package=pheatmap), with dendrograms based on Euclidean distance and the “ward.D” clustering method. When necessary, distances were imputed using the ultrametric function of the ape package^85, 86^.

### *Myo1d* Knock-down in Culture Mouse Embryos

E7.5 mouse embryos were harvested and cultured in DMEM with 50% Rat serum. Lenti-virus particles were added in medium during culture. Heart looping on *Myo1d* knockdown embryos were analyzed using a Leica M165FC microscope with ProgRes® C14 digital camera. E7.5 mouse embryos were harvested and cultured 5 hours in DMEM with 50% Rat serum at regular culture condition (37°C, 5% CO2) without rotation. Five hours later, *Myo1d* shRNA and Control shRNA lenti-virus particles were added in medium and embryos were cultured 48 hours with rotation. After 48 hours, embryos were fixed in 4% PFA/PBS. Heart looping on *Myo1d* knockdown embryos were analyzed using a Leica M165FC microscope with ProgRes® C14 digital camera.

*Myo1d* shRNA Lenti-virus particles were generated with RMM4431-200318212 (Dharmacon) by Genome Editing, Transgenic and Virus (GETV) Core at Magee-Womens Research Institute.

### *Myo1d* CRISPR/Cas9 Gene Edited Mice

C57BL/6NJ (JAX stock # 5304): and CB6F1/J mouse strains were used as embryo donors and pseudopregnant recipient dams, respectively. Cas9 (100 ng/µl) (TriLink Biotechnologies), gRNA (50 ng/µl each guide) were injected into the cytoplasm of zygotes. Oviduct transfers were performed on the same day into pseudopregnant dams. Founders were identified with the genotyping strategy described below, and the specific 334 bp deletion of exon 2 was confirmed in N1 progeny through TOPO TA cloning (Life Technologies) and Sanger sequencing. The resulting allele is designated *Myo1d*^em1(IMPC)J^.

Guides were designed against critical exon <u>ENSMUSE00001265867</u> of *Myo1d* using CRISPRtools (Peterson et al.) The following primers were used to generate PCR template for sgRNA synthesis as described^87^.

Myod1_u1:

5’-gaaattaatacgactcactataggACATTTGGTAATAAACTAAGgttttagagctagaaatagc-3’;

Myod1_u2:

5’-gaaattaatacgactcactatagGAAACAGTTACTAGGACATTgttttagagctagaaatagc-3’;

Myod1_d1:

5’-gaaattaatacgactcactatagGCTGTGATTCCAAAGCTGCAgttttagagctagaaatagc-3’;

Myod1_d2:

5’-gaaattaatacgactcactataggTTTTAAATACCTTGCAGCTTgttttagagctagaaatagc-3’.

PCR products were column purified (Qiagen), and *in vitro* transcription was performed with Ambion T7 MEGAshortscript kit (Life Technologies). RNA was recovered using an Ambion MEGAclear kit (Life Technologies) and final concentration was determined by NanoDrop (Thermo Scientific).

Genotyping of the gene targeted mice were conducted by PCR amplification using forward and reverse primers as follows: Exon2 F, 5’-GGAGGGAGAGAGATGGGATT-3’, Exon2 R, 5’-GCTTGGGCTTGACTTGAACT-3’, with the expected product sizes of 668 bp for wild-type and 344 bp for knock-out.

### Episcopic Confocal Microscopy Histopathology

E18.5 homozygote *Myo1d* mutant and control wildtype C57Bl/6 mice were embedded in paraffin for a detailed histopathology assessment of gut rotation using episcopic confocal microscopy (ECM). The serial image stacks were digitally resectioned in different imaging planes and viewed in 3D reconstructed to ascertain the precise position of the sigmoid colon relative to the superior mesenteric artery.

### Analysis of Gut Rotation in *myo1d* Zebrafish Mutants

Zebrafish were raised and kept under standard laboratory conditions at 28.5°C. To better visualize internal structures, embryos were incubated with 0.2 mM 1-phenyl-2-thiourea (Sigma) until 6dpf to inhibit pigmentation. Zebrafish *myo1d^pt31c^* (mutant was generated by TALENs as described^31^ in Saydmohammed et al., 2018. Gut orientation was analyzed by treating MZ *myo1d^pt31c^*and wildtype larvae with 0.3 μg/ml PED-6 (ThermoFisher #D23739) for 3 hours at 6 days post fertilization as described^88^, and scored for gut looping using epifluorescence microscopy (Leica M205 FA, IL). Survival data were plotted after counting settling down embryos over 7 days in larval rearing system. Survival curve was generated by using chi-square test. Comparison of survival curve were done by Mantel-Cox test for significance using Graphpad prism^89^.

### ToppFun/ToppCluster Network Analysis

To evaluate functional relationships of differentially expressed genes for each signature, we first used the ToppGene Suite/ToppFun tool, evaluating the enrichments table for each list, and then carried out a deeper global multi-feature network analysis using ToppCluster.cchmc.org using Gene Ontology, Mouse Phenotype, Pathways, protein interactions, conserved domains and TFBS annotations options. The entire results were downloaded from ToppGene and clustered semantically based on abstracted annotation terms shown and rendered using Morpheus for Feature Enrichments of Early/Late and Left/Right DEG genesets (-log10 p-value heatmap output).

### Human Heterotaxy Patient Analysis

Heterotaxy patients from the Children’s Hospital of Pittsburgh (n=67) and Children’s National Medical Center (CNMC) (n=6) were recruited after consenting under approved human study protocols approved by the University of Pittsburgh and CNMC Institutional Review Board. Blood was obtained for DNA extraction. Diagnosis and imaging data were obtained by review of the medical records.

Genomic DNA were processed for exome sequencing using Agilent V4 or V5 Exome Capture kits followed by sequencing using Illumina HiSeq2000/4000. Sequence alignment and variant calling were carried out using standard pipeline with BWA, Picard tools and GATK. VCF files were further annotated using Variant Effect Predictor (v89)^90^ with a single consequence retrieved for each variant. Analysis was conducted on the intersection of the Agilent V4 and V5 capture intervals using BEDTools (v2.26.0)^20^ and further filtered for variants with a genotyping quality >= 20 and read depth >= 20. High impact rare coding variants were defined as those coding variants with a gnomAD^91^ allele frequency <= 0.01 and a CADD^92^ PHRED score of at least 10. Mutations in patients HTX-9 and HTX-22 were validated by Sanger sequencing. For controls, we used exome sequencing data from 1000 Genomes 95 ancestry-matched CEU samples using the same intersection of the Agilent V4 and V5 capture intervals^93^[1000 Genomes Project, http://ftp.1000genomes.ebi.ac.uk/vol1/ftp/release/20130502/].

## URLs

DAVID Bioinformatics Resources 6.7, http://david.abcc.ncifcrf.gov/.

REVIGO, http://revigo.irb.hr/

ToppFun, https://toppgene.cchmc.org/enrichment.jsp

ToppCluster, https://toppcluster.cchmc.org

Morpheus, https://software.broadinstitute.org/morpheus/

## Data Availability

Data for *MYO1D* mutation is publicly available at the dbGAP under phs001691.v1.p1. The other data that support the findings of this study are available from the corresponding author upon reasonable request.

## Supporting information

All supplemental files

## ACKNOWLEDGEMENTS

This work was supported by funding from the AHA grant 847524 (H.Y), NIH grants HL098180 (C.W.L), HL157103 (C.W.L) and GM104412 (M.T, C.W.L).

## AUTHOR CONTRIBUTIONS

B.A contributed RNAseq bioinformatics and pathway enrichment analysis; C.B contributed Mouse *Myo1d* CRISPR gene editing; A.B contributed RNAseq bioinformatic data processing and analysis; C.C contributed Mouse embryo *Myo1d* shRNA analysis; W.D contributed Myo1d mutant mice gut looping phenotyping; G.G contributed Episcopic confocal microscopy for *Myo1D* KO mouse analysis; D.K contributed Bioinformatic analysis of bulk RNAseq and single cell RNAseq data; J.L contributed Analysis of heterotaxy patient clinical phenotypes; C.W.L contributed Study design, analysis of data, and manuscript preparation; M.M contributed Heterotaxy patient clinical phenotype determination; S.M contributed Construction of CRISPR gene edited *Myo1d* knockout mice; M.S contributed Construction of *myo1d* maternal zygotic zebrafish mutants, analysis for gut looping defects and impact on viability; M.T contributed Design of myo1d knockout zebrafish, analysis for gut looping defects, publication figures and manuscript preparation; H.Y contributed Study design, conduct RNAseq of mouse embryos, analysis of RNAseq data, comparisons with published single cell RNAseq data, Mouse embryo *Myo1d* shRNA analysis, *Myo1d* mutant mice gut looping phenotyping, publication figures and manuscript preparation; Y.W contributed MRI scans for phenotyping gut looping orientation in *Myo1d* knockout mice.

