## Supplementary material for "Spatial transcriptome profiling uncovers metabolic regulation of left-right patterning": All supplemental files

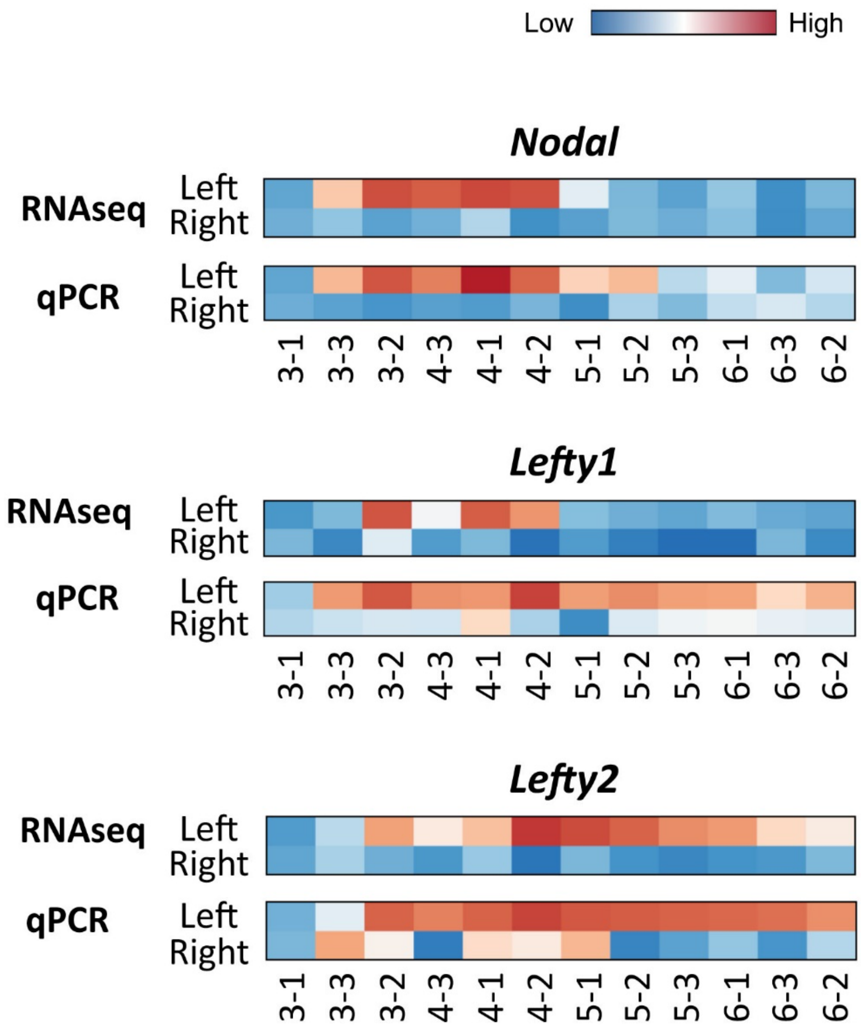


**Supplementary Figure 1. Real time PCR analysis of left and right half-embryos**

Efficacy of the microdissection of left and right half-embryos was demonstrated with real time PCR quantitation for *Nodal, Lefty1 and Lefty2* transcripts. Similar results were observed with qPCR as in the RNAseq data, but the qPCR showed higher detection sensitivity.

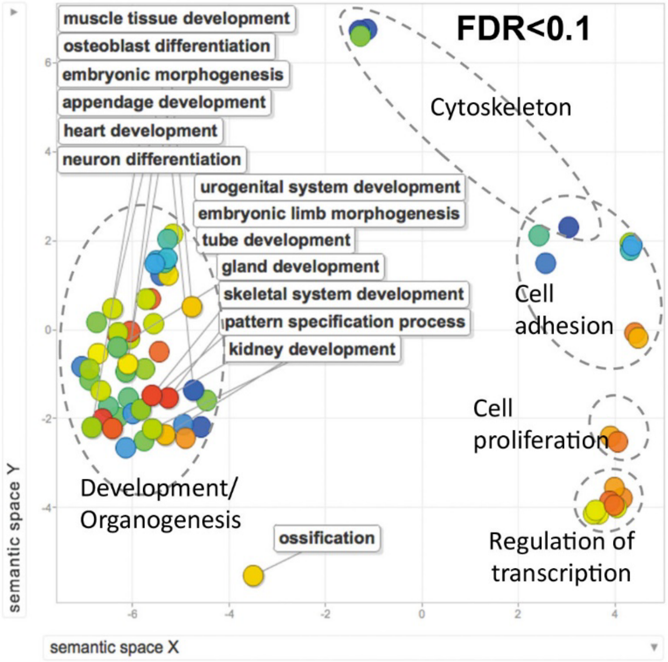


**Supplementary Figure 2. Gene ontology analysis of DEGs between the 3 and 6 somite stages**

DEGs recovered from the 3 to 6 somite stages were examined for pathway enrichment using DAVID[82], and visualized using Revigo[83]. This recovered an abundance of terms related to development.

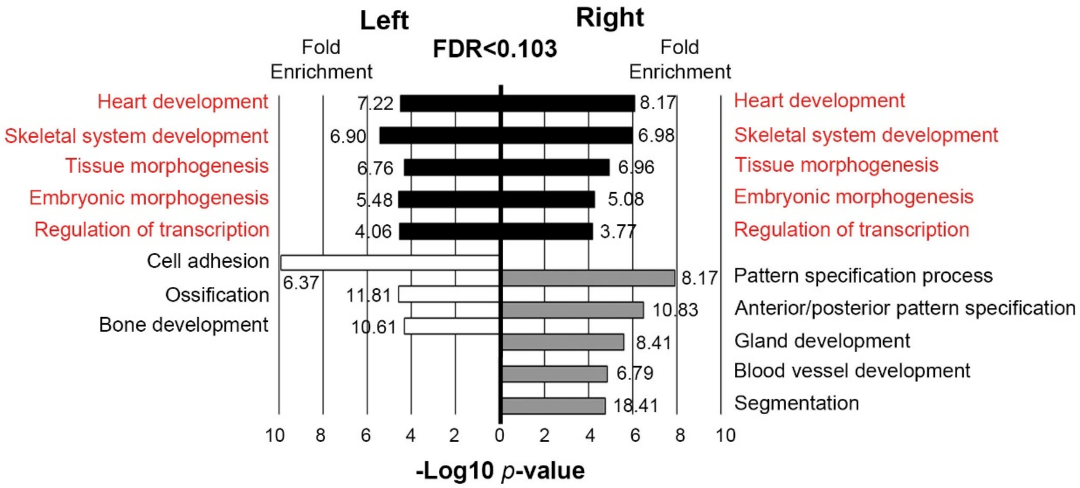


**Supplementary Figure 3. DEGs in the left and right half-embryos.**

DEGs in the left and right half-embryos from somite stages 3, 4, 5, and 6 were analyzed using GO Ontology. Many of the same pathways were recovered from the right and left half-embryos (indicated in red).

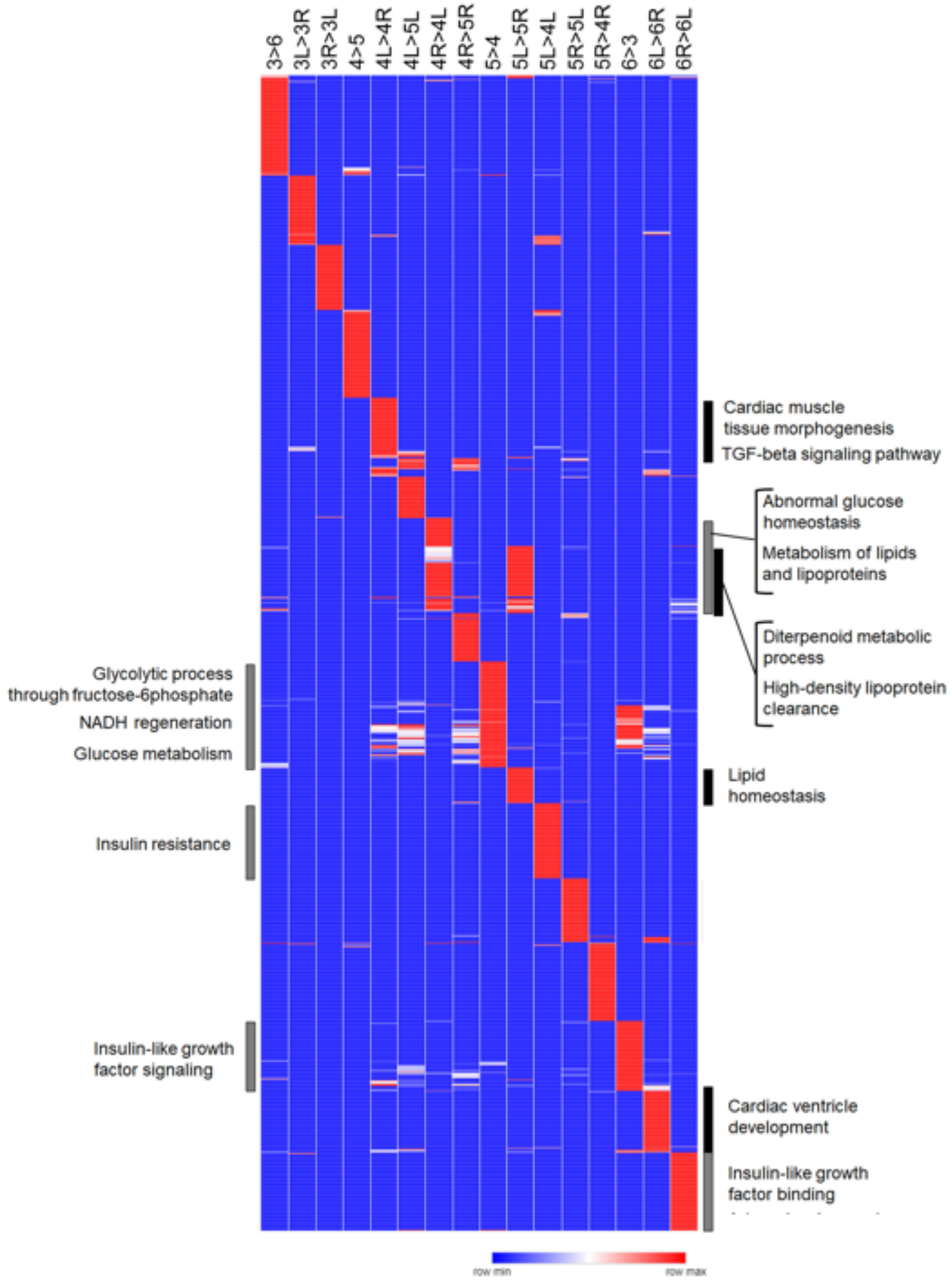


**Supplementary Figure 4. Heatmap of ToppCluster analysis of the RNASeq dataset**

ToppCluster analysis[94] of the entire RNAseq dataset was conducted and the heatmap generated using Morpheus (Broad Institute) is shown, with pathways shown recovered from comparisons between different developmental stages or between left vs. right half-embryos (see column headings). Heat map of DEGs more highly expressed, (>) or expressed at lower levels between sides or stages are shown, as denoted by the column headings. Notable from comparisons between stages and between left vs. right are many pathways related to insulin/IGF signaling and energy metabolism. Also notable are cardiac-related terms from the left vs. right comparisons, suggesting emergence of LR asymmetry in the embryonic heart. Left-sided TGFβ signaling was observed and in addition, the novel finding of left-sided Wnt signaling.

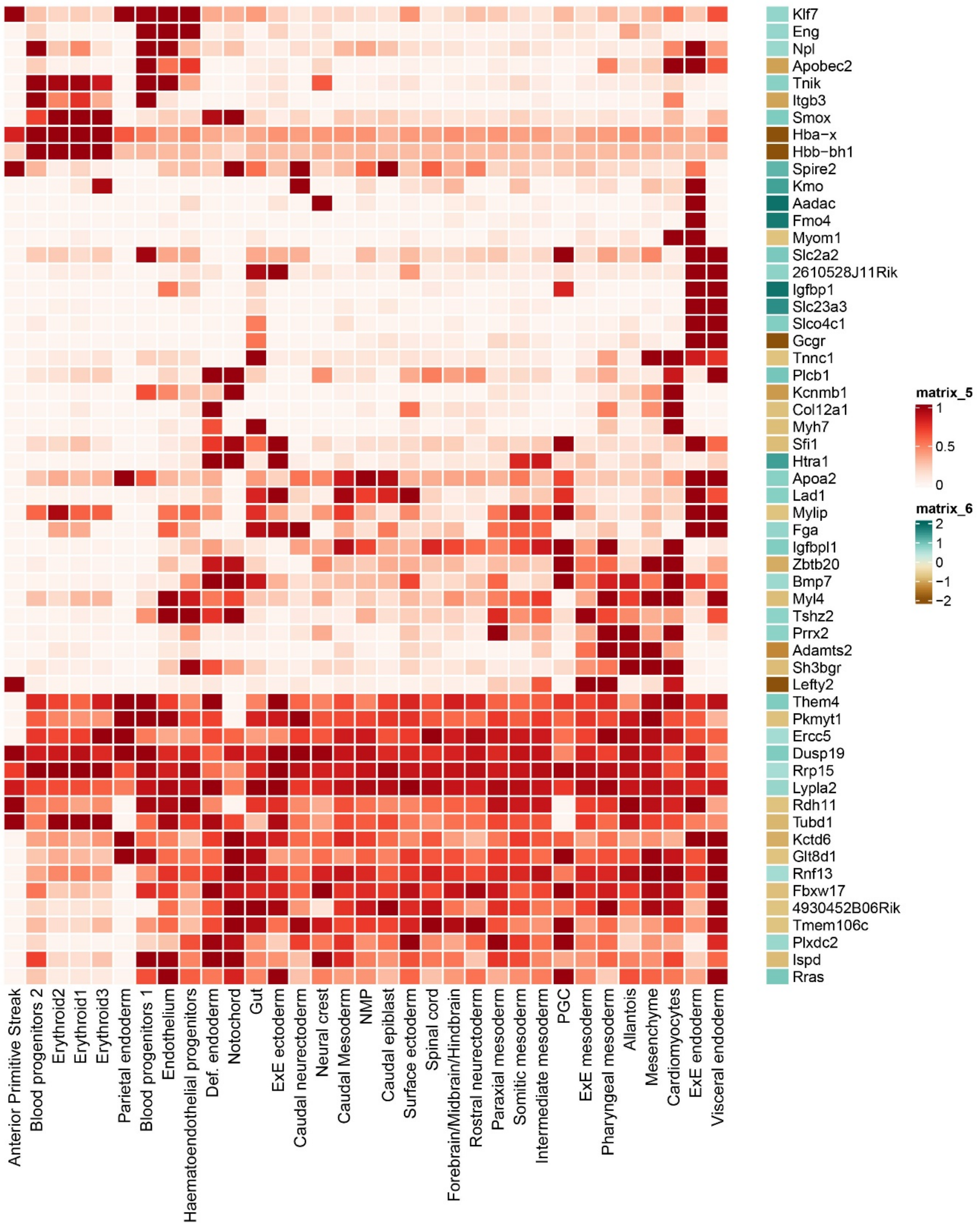


**Supplementary Figure 5. Expression of genes with high LR fold change in different lineages and cell types of the early mouse embryo.**

LR-DEGs identified in our spatial transcriptomic analysis of LR patterning was examined for their expression in different lineages or cell types using published single cell RNAseq data[40] at the equivalent 6S stage.

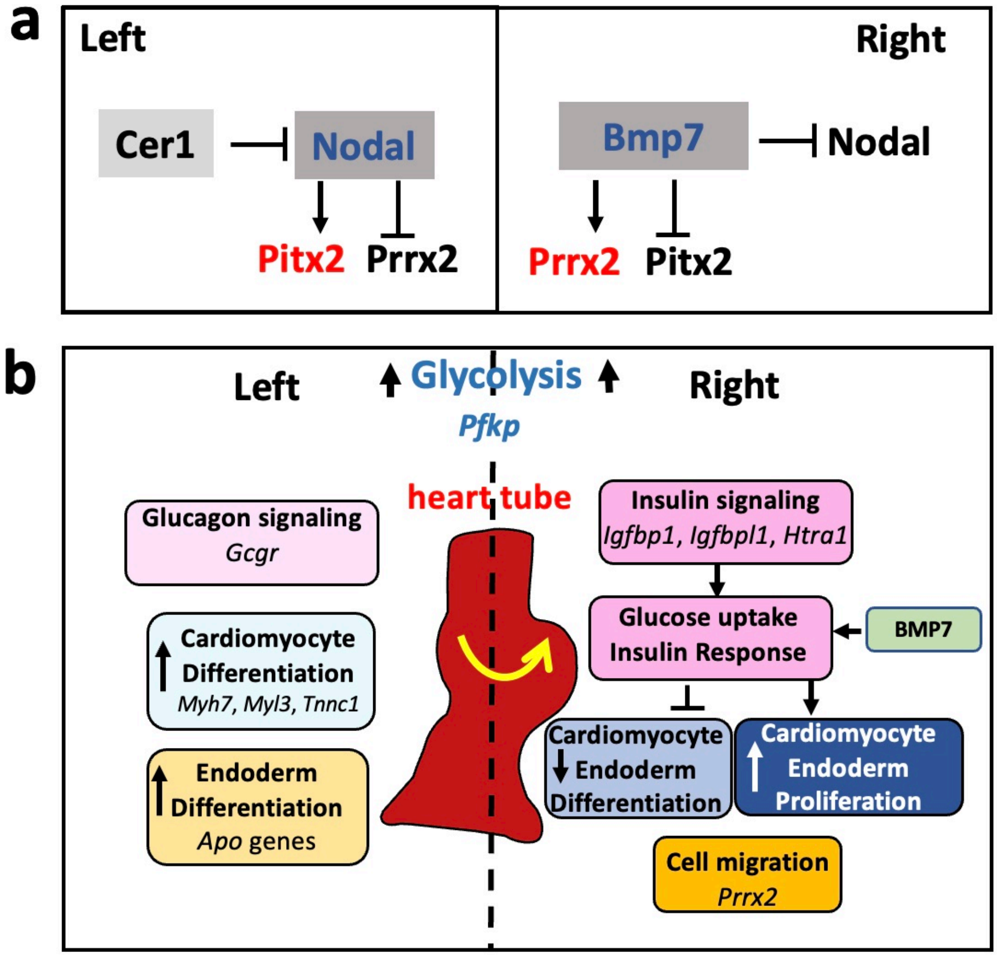


**Supplementary Figure 6. Schematic summary of cell signaling pathways and metabolic regulation of LR patterning**

**(a) Two parallel pathways establish the LR axis.**

To establish the LR body axis, left-sided activation of Nodal and right-sided BMP signaling are accompanied by downstream activation of the homeobox transcription factors *Pitx2* on the left, and *Prrx2* on the right. Nodal signaling, the key cell signaling pathway on the left, may undergo feedback regulation with left-sided *Cerl1* mediated repression and right sided silencing with *Bmp7* expression.

**(b). Metabolic regulation of LR patterning in the early mouse embryo.**

Higher IGF signaling on the right predicts increased proliferation and repressed differentiation of mesodermally derived cardiomyocytes and endodermally derived gut-liver cell lineages. Combined with enhanced EMT and cell migration mediated by right-sided *Prrx2*, this may promote right sided heart looping and contribute to patterning of liver/gut laterality.

**Table S1. *MYO1D* and Laterality gene mutations identified in heterotaxy patients**

| Patient. ID | Sex | Gene | Nucleotide Change | Amino Acid Change | CADD_PHRED | dbSNP | Allele Frequency gnomAD* | Transmission |
|---|---|---|---|---|---|---|---|---|
| 7007 | M | MYO1D | c.2312G>A | p.Arg771His | 20.3 | rs7215958 | 7.10E-03 | Father/cis |
| | | MYO1D | c.2293C>T | p.Pro765Ser | 23.1 | rs7209106 | 2.43E-03 | Father/cis |
| 7101 | F | MYO1D | c.2950C>T | p.Arg984Trp | 34 | NA | 4.49E-05 | Mother/cis |
| | | MYO1D | c.2585A>T | p.His862Leu | 21.5 | rs138039699 | 2.54E-03 | Mother/cis |

* Allele frequencies retrieved from Genome Aggregation Database (exome)
